# No evidence of human genome integration of SARS-CoV-2 found by long-read DNA sequencing

**DOI:** 10.1101/2021.05.28.446065

**Authors:** Nathan Smits, Jay Rasmussen, Gabriela O. Bodea, Alberto A. Amarilla, Patricia Gerdes, Francisco J. Sanchez-Luque, Prabha Ajjikuttira, Naphak Modhiran, Benjamin Liang, Jamila Faivre, Ira W. Deveson, Alexander A. Khromykh, Daniel Watterson, Adam D. Ewing, Geoffrey J. Faulkner

## Abstract

A recent study proposed severe acute respiratory syndrome coronavirus 2 (SARS-CoV-2) hijacks the LINE-1 (L1) retrotransposition machinery to integrate into the DNA of infected cells. If confirmed, this finding could have significant clinical implications. Here, we applied deep (>50×) long-read Oxford Nanopore Technologies (ONT) sequencing to HEK293T cells infected with SARS-CoV-2, and did not find the virus integrated into the genome. By examining ONT data from separate HEK293T cultivars, we completely resolved 78 L1 insertions arising *in vitro* in the absence of L1 overexpression systems. ONT sequencing applied to hepatitis B virus (HBV) positive liver cancer tissues located a single HBV insertion. These experiments demonstrate reliable resolution of retrotransposon and exogenous virus insertions via ONT sequencing. That we found no evidence of SARS-CoV-2 integration suggests such events are, at most, extremely rare *in vivo*, and therefore are unlikely to drive oncogenesis or explain post-recovery detection of the virus.

## INTRODUCTION

Severe acute respiratory syndrome coronavirus 2 (SARS-CoV-2) is a positive-sense single-stranded ∼30kbp polyadenylated RNA betacoronavirus (V’kovski et al., 2020; Wu et al., 2020). SARS-CoV-2 does not encode a reverse transcriptase (RT) and therefore is not expected to integrate into genomic DNA as part of its life cycle. This assumption is of fundamental importance to the accurate diagnosis and potential long-term clinical consequences of SARS-CoV-2 infection, as demonstrated by other viruses known to incorporate into genomic DNA, such as human immunodeficiency virus 1 (HIV-1) and hepatitis B virus (HBV) (Bill and Summers, 2004; Fujimoto et al., 2012; Jiang et al., 2012; Nagaya et al., 1987). Notably, a recent work by Zhang *et al*. reported potential evidence of SARS-CoV-2 integration into the genome of infected human cells (Zhang et al., 2021). Prior analyses of mammalian genome sequences, as well as *in vivo* and *in vitro* experimental data, indicate single-stranded RNA viruses can act as templates for endogenous RTs (Belyi et al., 2010; Feschotte and Gilbert, 2012; Geuking et al., 2009; Horie et al., 2010; Kawasaki et al., 2021; Klenerman et al., 1997). These studies provide a conceptual basis to further investigate genomic integration of SARS-CoV-2, as pursued by Zhang *et al*..

LINE-1 (L1) retrotransposons reside in all mammalian genomes (Kazazian and Moran, 2017). In humans, L1 transcribes a bicistronic mRNA encoding two proteins, ORF1p and ORF2p, essential to L1 mobility (Moran et al., 1996). ORF2p possesses endonuclease (EN) and RT activities, and exhibits strong *cis* preference for reverse transcription of L1 mRNA (Doucet et al., 2015; Garcia-Perez et al., 2007; Kulpa and Moran, 2006; Monot et al., 2013; Moran et al., 1996; Wei et al., 2001). Nonetheless, the L1 protein machinery can *trans* mobilise polyadenylated cellular RNAs, especially those produced by *Alu* and SINE-VNTR-*Alu* (SVA) retrotransposons, but also including protein-coding gene mRNAs (Dewannieux et al., 2003; Esnault et al., 2000; Garcia-Perez et al., 2007; Hancks et al., 2011; Raiz et al., 2012). Somatic L1 mobilisation *in cis* is observed in embryonic cells, the neuronal lineage, and various cancers (Evrony et al., 2015; Feusier et al., 2019; Rodriguez-Martin et al., 2020; Sanchez-Luque et al., 2019; Schauer et al., 2018; Scott et al., 2016). By contrast, somatic L1-mediated *trans* mobilisation is apparently rare *in vivo* (Evrony et al., 2015; Rodriguez-Martin et al., 2020; Sanchez-Luque et al., 2019) and is likely repressed by various mechanisms (Ahl et al., 2015; Deniz et al., 2019; Doucet et al., 2015; Ewing et al., 2020; Sanchez-Luque et al., 2019). While *Alu* and, to a lesser extent, SVA are readily mobilised in cultured cell assays by L1 proteins, the same machinery produces less than one non-retrotransposon cellular RNA insertion for every 2000 L1 insertions (Dewannieux et al., 2003; Hancks et al., 2011; Wei et al., 2001). Both *cis* and *trans* L1-mediated insertions incorporate target site duplications (TSDs) and a 3′ polyA tract, and integrate at the degenerate L1 EN motif 5′-TTTT/AA (Dewannieux et al., 2003; Esnault et al., 2000; Garcia-Perez et al., 2007; Gilbert et al., 2005; Hancks et al., 2009; Jurka, 1997; Moran et al., 1996; Raiz et al., 2012). These sequence hallmarks can together discriminate artifacts from genuine insertions (Faulkner and Billon, 2018).

In their work, Zhang *et al*. overexpressed L1 in HEK293T cells, infected these with SARS-CoV-2, and identified DNA fragments of the virus through PCR amplification. These results, alongside other less direct (Kazachenka and Kassiotis, 2021; Yan et al., 2021) analyses, were interpreted as evidence of SARS-CoV-2 genomic integration. Crucially, Zhang *et al*. then detected 63 putative SARS-CoV-2 integrants by Oxford Nanopore Technologies (ONT) long-read sequencing. Of these, only a single integrant on chromosome X was spanned by an ONT read aligned to one locus, and was flanked by potential TSDs (**Figure 1A**). However, this SARS-CoV-2 integrant did not incorporate a 3′ polyA tract, as is expected for an L1-mediated insertion, and involved an unusual 28kb internal deletion of the SARS-CoV-2 sequence. Overall, the SARS-CoV-2 integrants reported by Zhang *et al*. were 26-fold enriched in exons, despite the L1 EN showing no preference for these regions (Flasch et al., 2019; Sultana et al., 2019). Zhang *et al*. also used Illumina short-read sequencing to map putative SARS-CoV-2 integration junctions in HEK293T cells without L1 overexpression. A lack of spanning reads and the tendency of Illumina library preparation to produce artefacts (Treiber and Waddell, 2017) leave this analysis open to interpretation.

**Figure 1.**
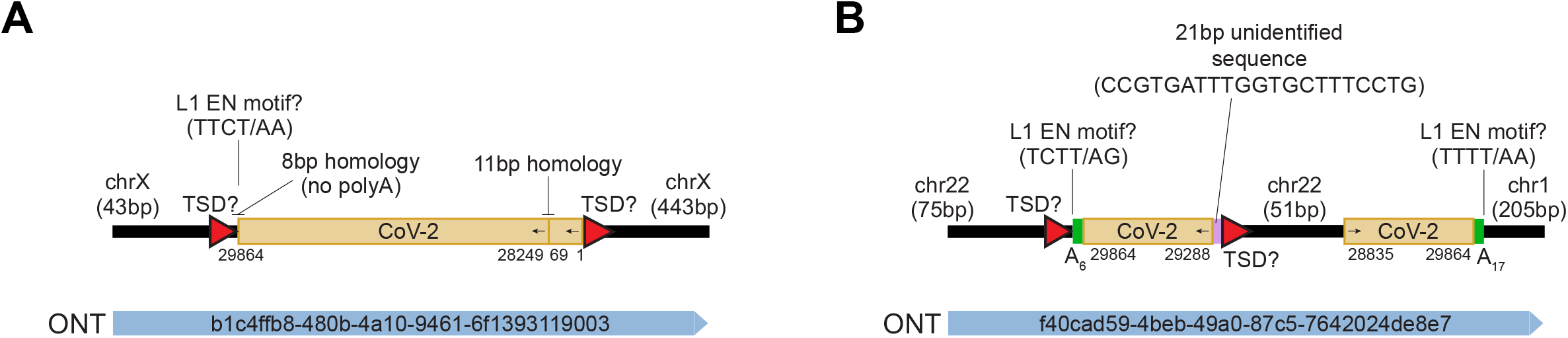
Key potential SARS-CoV-2 insertions reported by Zhang *et al*.. **(A)** A cartoon summarising the features of a putative SARS-CoV-2 integrant on chromosome X. Numerals underneath the SARS-CoV-2 sequence represent positions relative to the QLD02 virus isolate. Potential TSDs are shown as red triangles, and motifs resembling potential pre-integration L1 EN recognition sites are highlighted, with question marks in labels intended to flag uncertain L1 involvement. No 3’ polyA tract was found. Homologous regions at sequence junctions are marked. One spanning ONT read is positioned underneath the cartoon and its identifier is displayed. **(B)** As for (A), except showing an ONT read spanning two SARS-CoV-2 insertions, on chromosome 22 and chromosome 1. The alignments to chromosome 22 were flagged as supplementary by the minimap2 aligner. 3’ polyA tracts are represented as green rectangles. Note: these chromosome 22 and chromosome X instances are the key examples reported by Zhang *et al*. in support of SARS-CoV-2 genomic integration. Neither example has a complete set of retrotransposition hallmarks (TSD, 3’ polyA tract, L1 EN motif) *and* the support of a uniquely aligned ONT read.

The application of ONT sequencing to HEK293T cells nonetheless held conceptual merit. ONT reads can span germline and somatic retrotransposition events end-to-end, and resolve the sequence hallmarks of L1-mediated integration (Ewing et al., 2020; Siudeja et al., 2021). Through this approach, we previously found two somatic L1 insertions in the liver tumour sample of an individual positive for hepatitis C virus (HCV), a ∼10kbp positive-sense single-stranded non-polyadenylated RNA virus (Lauer and Walker, 2001), including one PCR-validated L1 insertion spanned by a single ONT read (Ewing et al., 2020; Shukla et al., 2013). Moreover, HEK293T cells are arguably a context favourable to L1 activity. They express L1 ORF1p (Philippe et al., 2016), accommodate engineered L1-mediated retrotransposition *in cis* and *in trans* (Hancks et al., 2011; Kubo et al., 2006; Niewiadomska et al., 2007; Sanchez-Luque et al., 2019), and support SARS-CoV-2 viral replication (**Figure S1**). Endogenous L1-mediated insertions can be detected in cell culture by genomic analysis of separate cultivars derived from a common population (Klawitter et al., 2016; Nguyen et al., 2018). Based on this experimental rationale, we sought to replicate the central findings of Zhang *et al*. and, after deeply ONT sequencing SARS-CoV-2-infected HEK293T cells, did not detect SARS-CoV-2 genomic integration.

## RESULTS

We ONT sequenced (∼54× genome-wide depth, read length N50 ∼ 39kbp) genomic DNA harvested from HEK293T cells infected with SARS-CoV-2 at a multiplicity of infection (MOI) of 1.0, as well as mock infected cells (∼28× depth, N50 ∼ 47kbp) (**Figures 2A** and **S1**, and **Table S1**). As positive controls, we ONT sequenced the tumour and non-tumour liver tissue of a HBV-positive hepatocellular carcinoma patient (Shukla et al., 2013). HBV is a DNA virus known to be integrated into sites of genomic damage via DNA double-strand break repair (Bill and Summers, 2004). As negative controls, we used the aforementioned HCV-positive hepatocellular carcinoma samples, and a normal liver sample (Ewing et al., 2020) (**Table S1**). We viewed HCV-infected samples as a suitable negative control because HCV and SARS-CoV-2 are both positive-sense single-stranded RNA viruses, yet HCV is not polyadenylated and is therefore unlikely to attract the L1 machinery, and has not been found to integrate into infected hepatocytes or liver tumour genomes (Fujimoto et al., 2012; Lauer and Walker, 2001). To these data, we added those of Zhang *et al*., and then used the Transposons from Long DNA Reads (TLDR) (Ewing et al., 2020) software to call SARS-CoV-2, HBV, HCV and non-reference retrotransposon insertions spanned by at least one uniquely aligned ONT read. TLDR detected no SARS-CoV-2, HBV or HCV insertions.

**Figure 2.**
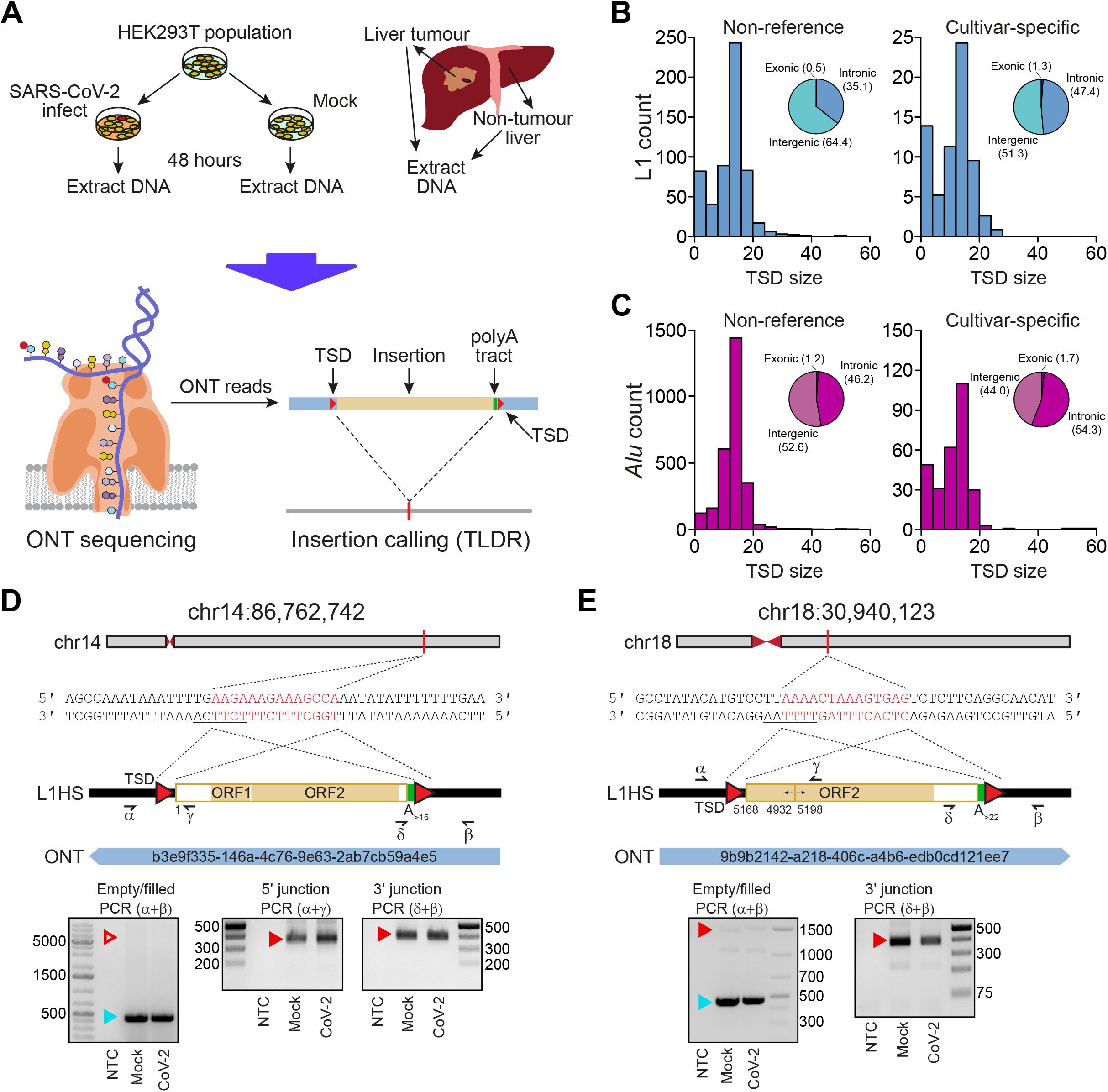
Detection of endogenous L1-mediated retrotransposition in human cells. **(A)** Experimental design. HEK293T cells were divided into two populations (cultivars), which were then either SARS-CoV-2 infected or mock infected. DNA was extracted from each cultivar, as well as from hepatocellular carcinoma patient samples, and subjected to ONT sequencing. ONT reads were used to call non-reference L1 and virus insertions with TLDR, which also resolves TSDs and other retrotransposition hallmarks. TSDs: red triangles; polyA tract: green rectangle; ONT read: blue rectangle. Some illustrations are adapted from a previous study (Ewing et al., 2020). **(B)** TSD size distribution for non-reference L1 insertions, as annotated by TLDR, as well as cultivar-specific L1 insertions found only in either our HEK293T cells infected with SARS-CoV-2 or our mock infected cells. Pie charts indicate the percentages of exonic, intronic and intergenic insertions, annotated by RefSeq coordinates. **(C)** As for (B), except showing data for *Alu* insertions. **(D)** Detailed characterisation of an L1 insertion detected in SARS-CoV-2 infected HEK293T cells by a single spanning ONT read aligned to chromosome 14. Nucleotides highlighted in red correspond to the integration site TSD. Underlined nucleotides correspond to the L1 EN motif. The cartoon indicates a full-length L1HS insertion flanked by TSDs (red triangles), and a 3’ polyA tract (green), with the underneath numeral representing the 5’ end position relative to the mobile L1HS sequence L1.3 (Dombroski et al., 1993). The relevant spanning ONT read, with identifier, is also positioned underneath the cartoon. Symbols (α, β, δ, γ) represent the approximate position of primers used for empty/filled site and L1-genome junction PCR validation reactions. These are displayed in gel images if successful. Ladder band sizes are as indicated, NTC; non-template control. Red triangles indicate L1 amplicon expected sizes (empty triangle: no product; filled triangle: capillary sequenced on-target product). Blue triangles indicate expected empty site sizes. **(E)** As for (D), except for a 5’ inverted/deleted L1HS located on chromosome 18. Please see Figures S1 and S2, and Tables S1 and S2 for further information.

In total, TLDR identified 575 non-reference human-specific L1 (L1HS) insertions, which were typically flanked by TSDs with a median length of 14bp (**Figure 2B** and **Table S2**). No tumour-specific L1 insertions were found, apart from the two previously detected in the HCV-infected liver tumour (Ewing et al., 2020; Shukla et al., 2013). Seventy-eight L1 insertions were found only in our SARS-CoV-2 infected HEK293T cells (66) or the mock infected control (12) and produced TSDs with a median length of 14bp (**Figure 2B**). Of the 78 events, 69 (88.5%) were detected by a single spanning read and 13 carried a 3′ transduction (Holmes et al., 1994; Moran et al., 1999) (**Table S2**). After random downsampling, the more deeply sequenced SARS-CoV-2 infected HEK293T cells still had more than 2-fold more putative cultivar-specific L1 insertions than the mock infected HEK293T cells. Next, we chose at random 6/69 L1 insertions detected by one spanning read for manual curation and PCR validation. All 6 L1 insertions bore a TSD and a 3′ polyA tract, and integrated at a degenerate L1 EN motif (**Figures 2D, 2E** and **S2A-S2D**). Three were 5′ inverted (Kazazian et al., 1988; Ostertag and Kazazian, 2001) (**Figures 2E, S2C** and **S2D**) and one carried a 3′ transduction (Holmes et al., 1994) traced to a mobile (Rodriguez-Martin et al., 2020) full-length non-reference L1HS (**Figure S2C**). Three PCR amplified in the SARS-CoV-2 and mock infected samples (**Figures 2D, 2E** and **S2A**) and three did not amplify in either sample (**Figures S2B-S2D**). The 6 integration sites were on average spanned by 86 reads not containing the L1 insertion (**Figure S2E**), a ratio (1:86) suggesting the L1s were absent from most cells. An additional analysis revealed 293 putative *Alu* (291) and SVA (2) insertions specific to either one of the HEK293T populations, with 290 of these found in the SARS-CoV-2 infected cells and 275 (93.9%) detected by a single spanning read (**Table S2**). The median TSD size for this cohort was 13bp (**Figure 2C**). Altogether, these and earlier (Ewing et al., 2020; Siudeja et al., 2021) experiments show that lone spanning ONT reads can recover *bona fide* retrotransposition events, and highlight endogenous L1 activity in HEK293T cells lacking L1 overexpression systems.

We next tested whether our computational analysis parameters excluded genuine HBV, HCV or SARS-CoV-2 insertions. We directly aligned our ONT reads to the genome of the SARS-CoV-2 isolate (QLD002, GISAID EPI_ISL_407896) used here, as well as to a geographically diverse set of HBV and HCV genomes (**Table S1**), and a highly mobile L1HS sequence (Dombroski et al., 1993). In total, 3.6% of our ONT sequence bases aligned to L1HS, whereas no alignments to the SARS-CoV-2 or HCV genomes were observed (**Figure 3A**). One read from the HBV-infected non-tumour liver sample aligned to 2,770bp of a HBV genotype B isolate, and the remaining 2,901bp aligned to an intergenic region of chromosome 2 (**Figure 3B** and **Table S2**). To validate this HBV insertion, we PCR amplified and capillary sequenced its 3′ junction (**Figure 3B**). The HBV sequence was linearised and rearranged (**Figure 3B**) as per prior reports (Fujimoto et al., 2012; Jiang et al., 2012; Nagaya et al., 1987). Direct inspection of ONT read alignments thus recovered a HBV integrant, which are found in ∼1 per 10_1_-10_4_ infected hepatocytes (Mason et al., 2016; Tu et al., 2018), yet did not reveal reads alignable to the SARS-CoV-2 genome in our ONT datasets.

**Figure 3.**
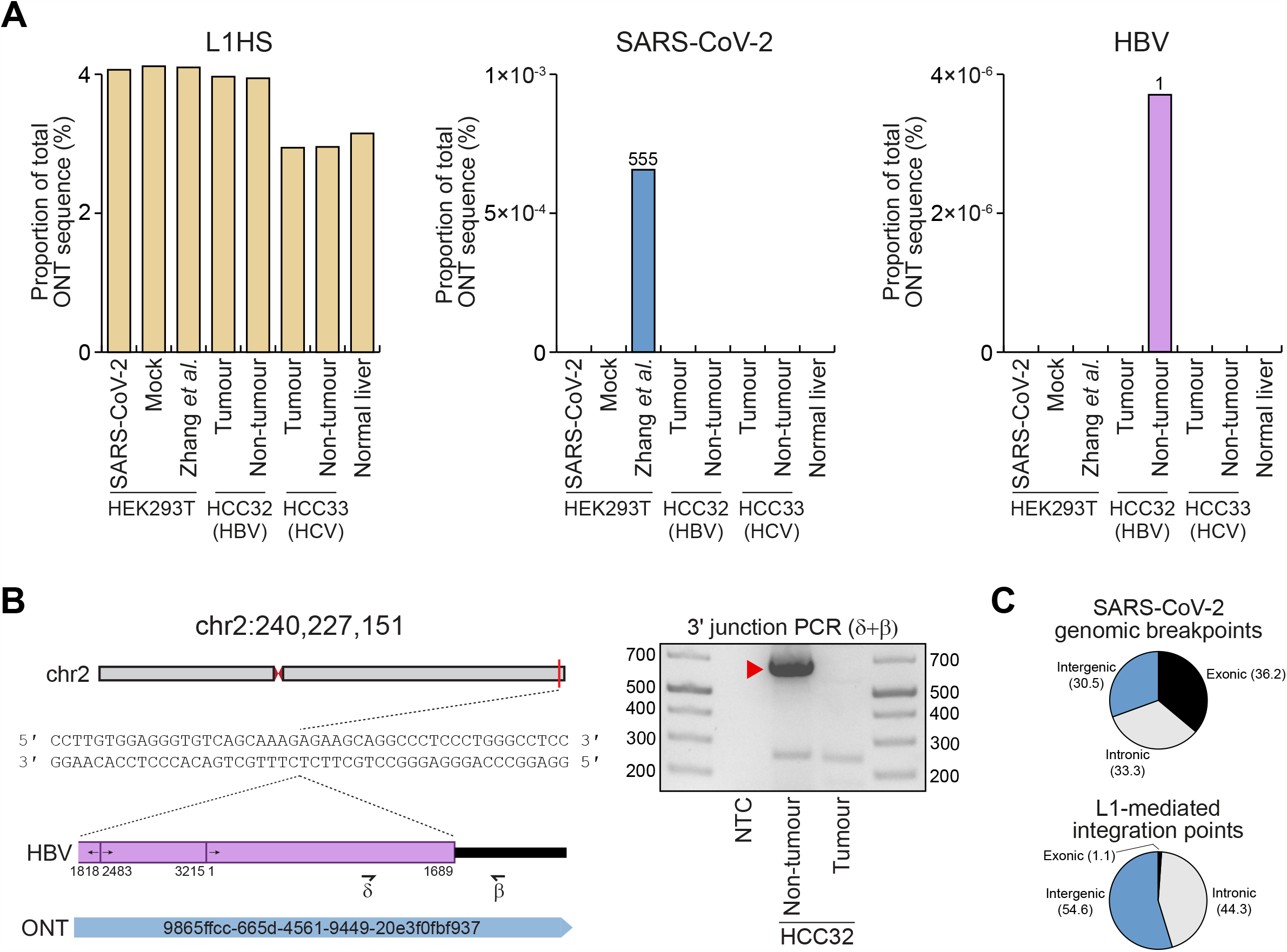
ONT reads occasionally align to viral genome sequences. **(A)** Percentages of total ONT sequence alignable to L1HS (left), SARS-CoV-2 (middle) and HBV (right) isolate genomes. Read counts for SARS-CoV-2 and HBV are provided above histogram columns. No reads were aligned to the HCV isolate genomes. HEK293T data were generated here (SARS-CoV-2, mock) or by Zhang *et al*.. HCC tumour/non-tumour liver pairs were sequenced here (HCC32; confirmed HBV-positive) or previously (Ewing et al., 2020) (HCC33; HCV-positive). Normal liver ONT sequencing from our prior work (Ewing et al., 2020) was included as an additional control. **(B)** A HBV insertion detected in non-tumour liver. In this example, an ONT read from the non-tumour liver of HCC32 spanned the 3’ junction of a HBV integrant located on chromosome 2. Of the HBV isolate genomes considered here, this read aligned best to a representative of genotype B (Genbank accession AB602818). The HBV sequence was rearranged consistent with its linearisation prior to integration (Fujimoto et al., 2012; Jiang et al., 2012; Nagaya et al., 1987). Numerals indicate positions relative to AB602818. Symbols (β, δ) represent the approximate position of primers used to PCR validate the HBV insertion. The gel image at right shows the 3’ junction PCR results. Ladder band sizes are as indicated. The red filled triangle indicates an on-target product confirmed by capillary sequencing. Repeated attempts to PCR amplify the 5’ junction of the HBV integrant did not return an on-target product, perhaps due to a genomic deletion at the insertion site. **(C)** Percentages of exonic (black), intronic (grey) and intergenic (blue) genomic alignment breakpoints for ONT reads also aligned to the SARS-CoV-2 genome, and also for the non-reference L1-mediated insertions reported here. Genomic features were annotated according to RefSeq coordinates. Please see Tables S1 and S2 for further information.

Reanalysing the ONT data generated by Zhang *et al*., we found 555 reads (out of ∼12 million) that generated an alignment of ≥100bp to the SARS-CoV-2 genome (**Figure 3A**). These reads (median length 924bp) were however 65.6% shorter than the overall dataset (2,686kbp) and were comprised of a much higher average proportion of SARS-CoV-2 sequence (52.3%) than the proportion of L1HS sequence found in reads aligned to L1HS (17.1%). Of the 555 reads, 79 generated an alignment of ≥100bp to the human genome, including one matching the aforementioned integrant on chromosome X that lacked a 3′ polyA tract (**Figure 1A**). An analysis of the corresponding 79 human genome alignment breakpoints, which could be interpreted as putative SARS-CoV-2 insertion points, as per Zhang *et al*., indicated 36.2% were exonic (**Figure 3C**). By comparison, 1.1% of all non-reference L1-mediated insertions reported here were exonic (**Figure 3C**), as were 1.3% and 1.7% of cultivar-specific L1 and *Alu* insertions, respectively (**Figures 2B** and **2C**). Finally, we investigated why TLDR called neither of the two SARS-CoV-2 insertions highlighted by Zhang *et al*. (**Figure 1**), and found ambiguity in the ONT read alignments supporting these examples led to their exclusion. Specifically, the putative chromosome X insertion (**Figure 1A**) was filtered due to its short 5′ genomic flank alignment, whereas the chromosome 22 example was filtered because the corresponding ONT read alignment was marked as supplementary to another alignment on chromosome 1 (**Figure 1B**). These analyses confirmed SARS-CoV-2 alignable reads were present in the Zhang *et al*. ONT dataset, yet these reads were unusually short and could include molecular artifacts interpreted by Zhang *et al*. as SARS-CoV-2 integrants.

## DISCUSSION

We did not observe L1-mediated SARS-CoV-2 genomic integration in HEK293T cells, despite availability of the L1 machinery (Hancks et al., 2011; Kubo et al., 2006; Niewiadomska et al., 2007; Philippe et al., 2016; Sanchez-Luque et al., 2019) and detected L1, *Alu* and SVA retrotransposition events. The higher number of L1 and *Alu* insertions found in our SARS-CoV-2-infected HEK293T cells is of potential interest given viral infection can repress host factors limiting L1 activity (Hrecka et al., 2011; Laguette et al., 2011; Zhao et al., 2013). This preliminary finding perhaps indicates SARS-CoV-2 infection could increase L1 or *Alu* retrotransposition *in vitro*, a possibility requiring experimental confirmation. The comparative rarity of SVA insertions, and absence of SARS-CoV-2 insertions, is however congruent with the relative frequencies of L1, *Alu*, SVA and non-retrotransposon cellular RNA insertions driven by L1 proteins in prior cultured cell assays (Dewannieux et al., 2003; Hancks et al., 2011; Wei et al., 2001).

Our approach has several notable differences and caveats when compared to that of Zhang *et al*.. Each study used different SARS-CoV-2 isolates, and here the multiplicity of infection (MOI 1.0) was double that of Zhang *et al*. (MOI 0.5). The high molecular weight DNA extraction method, ONT library preparation kit and depth and quality of sequencing applied to HEK293T cells by Zhang *et al*. (standard isopropanol precipitation, SQK-LSK109 kit, ∼21× depth, N50 ∼ 11kbp) and here (Nanobind kit, SQK-LSK110 kit, ∼54× depth, N50 ∼ 39kbp) differed. Nevertheless, the DNA extraction protocols of each study would limit retention of extrachromosomal SARS-CoV-2 DNA potentially generated by ectopic L1 reverse transcription (Dhellin et al., 1997). The origins of the ONT reads aligned to the SARS-CoV-2 genome reported by Zhang *et al*. are therefore unclear in our view. Zhang *et al*. only ONT sequenced HEK293T cells transfected with an L1 expression plasmid, which human cells would not carry *in vivo*. We did not analyse SARS-CoV-2 patient samples although, arguably, HEK293T cells present an environment far more conducive to L1 activity than those cells accessed *in vivo* by SARS-CoV-2 (Sungnak et al., 2020; Wiersinga et al., 2020). Widespread cell death post-infection also reduces the probability SARS-CoV-2 integrants would persist in the body (Karki et al., 2021; Varga et al., 2020). This view aligns with a very recent report of negligible SARS-CoV-2 DNA being detected by PCR in COVID-19 patient nasal swabs (Briggs et al., 2021).

Finally, the incredible enrichment reported by Zhang *et al*. for putative SARS-CoV-2 insertions in exons, which this and prior studies (Flasch et al., 2019; Sultana et al., 2019) have shown are not preferred by the L1 EN, contradicts the involvement of L1 in the events interpreted by Zhang *et al*. as SARS-CoV-2 genomic integrants. We conclude L1 *cis* preference strongly disfavours SARS-CoV-2 retrotransposition, making the phenomenon mechanistically plausible but likely very rare, as for other polyadenylated non-retrotransposon cellular RNAs (Dewannieux et al., 2003; Doucet et al., 2015; Esnault et al., 2000; Garcia-Perez et al., 2007; Hancks et al., 2011; Kulpa and Moran, 2006; Monot et al., 2013; Moran et al., 1996; Wei et al., 2001).

## ACKNOWLEDGEMENTS

The authors thank the human subjects of this study who donated tissues to the Centre Hépatobiliaire, Paul-Brousse Hospital. We thank S. Richardson and R. Shukla for helpful discussions, K. Chappell and P. Young for project support, and Queensland Health for providing the SARS-CoV-2 virus isolate QLD02. This study was funded by an Australian Government Research Training Program (RTP) Scholarships (N.S.), an NHMRC-ARC Dementia Research Development Fellowship (GNT1108258, G.O.B.), seed funding provided by the Australian Infectious Disease Research Centre to establish SARS-CoV-2 research at the University of Queensland (A.A.K), an Australian Department of Health Medical Research Future Fund (MRFF) Novel Coronavirus Vaccine Development Grant (APP1202445-2020, D.W.), an MRFF Investigator Grant (MRF1175457, A.D.E.), an NHMRC Investigator Grant (GNT1173711, G.J.F.), a CSL Centenary Fellowship (G.J.F.), and the Mater Foundation.

## AUTHOR CONTRIBUTIONS

N.S., J.R., G.O.B., A.A.A., P.G., F.J.S-L., P.A., N.M. and B.L. performed experiments and analysed data. A.D.E. and G.J.F. performed bioinformatic analysis. J.F., I.W.D., A.A.K., D.W. and G.J.F. provided resources. G.J.F. designed the project and wrote the manuscript.

## FIGURE LEGENDS

**Figure 1. Key potential SARS-CoV-2 insertions reported by Zhang *et al*.**

**(A)** A cartoon summarising the features of a putative SARS-CoV-2 integrant on chromosome X. Numerals underneath the SARS-CoV-2 sequence represent positions relative to the QLD02 virus isolate. Potential TSDs are shown as red triangles, and motifs resembling potential pre-integration L1 EN recognition sites are highlighted, with question marks in labels intended to flag uncertain L1 involvement. No 3’ polyA tract was found. Homologous regions at sequence junctions are marked. One spanning ONT read is positioned underneath the cartoon and its identifier is displayed.

**(B)** As for (A), except showing an ONT read spanning two SARS-CoV-2 insertions, on chromosome 22 and chromosome 1. The alignments to chromosome 22 were flagged as supplementary by the minimap2 aligner. 3’ polyA tracts are represented as green rectangles. Note: these chromosome 22 and chromosome X instances are the key examples reported by Zhang *et al*. in support of SARS-CoV-2 genomic integration. Neither example has a complete set of retrotransposition hallmarks (TSD, 3’ polyA tract, L1 EN motif) *and* the support of a uniquely aligned ONT read.

**Figure 2. Detection of endogenous L1-mediated retrotransposition in human cells**.

**(A)** Experimental design. HEK293T cells were divided into two populations (cultivars), which were then either SARS-CoV-2 infected or mock infected. DNA was extracted from each cultivar, as well as from hepatocellular carcinoma patient samples, and subjected to ONT sequencing. ONT reads were used to call non-reference L1 and virus insertions with TLDR, which also resolves TSDs and other retrotransposition hallmarks. TSDs: red triangles; polyA tract: green rectangle; ONT read: blue rectangle. Some illustrations are adapted from a previous study (Ewing et al., 2020).

**(B)** TSD size distribution for non-reference L1 insertions, as annotated by TLDR, as well as cultivar-specific L1 insertions found only in either our HEK293T cells infected with SARS-CoV-2 or our mock infected cells. Pie charts indicate the percentages of exonic, intronic and intergenic insertions, annotated by RefSeq coordinates.

**(C)** As for (B), except showing data for *Alu* insertions.

**(D)** Detailed characterisation of an L1 insertion detected in SARS-CoV-2 infected HEK293T cells by a single spanning ONT read aligned to chromosome 14. Nucleotides highlighted in red correspond to the integration site TSD. Underlined nucleotides correspond to the L1 EN motif. The cartoon indicates a full-length L1HS insertion flanked by TSDs (red triangles), and a 3’ polyA tract (green), with the underneath numeral representing the 5’ end position relative to the mobile L1HS sequence L1.3 (Dombroski et al., 1993). The relevant spanning ONT read, with identifier, is also positioned underneath the cartoon. Symbols (α, β, δ, γ) represent the approximate position of primers used for empty/filled site and L1-genome junction PCR validation reactions. These are displayed in gel images if successful. Ladder band sizes are as indicated, NTC; non-template control. Red triangles indicate L1 amplicon expected sizes (empty triangle: no product; filled triangle: capillary sequenced on-target product). Blue triangles indicate expected empty site sizes.

**(E)** As for (D), except for a 5’ inverted/deleted L1HS located on chromosome 18. Please see Figures S1 and S2, and Tables S1 and S2 for further information.

**Figure 3. ONT reads occasionally align to viral genome sequences**.

**(A)** Percentages of total ONT sequence alignable to L1HS (left), SARS-CoV-2 (middle) and HBV (right) isolate genomes. Read counts for SARS-CoV-2 and HBV are provided above histogram columns. No reads were aligned to the HCV isolate genomes. HEK293T data were generated here (SARS-CoV-2, mock) or by Zhang *et al*.. HCC tumour/non-tumour liver pairs were sequenced here (HCC32; confirmed HBV-positive) or previously (Ewing et al., 2020) (HCC33; HCV-positive). Normal liver ONT sequencing from our prior work (Ewing et al., 2020) was included as an additional control.

**(B)** A HBV insertion detected in non-tumour liver. In this example, an ONT read from the non-tumour liver of HCC32 spanned the 3’ junction of a HBV integrant located on chromosome 2. Of the HBV isolate genomes considered here, this read aligned best to a representative of genotype B (Genbank accession AB602818). The HBV sequence was rearranged consistent with its linearisation prior to integration (Fujimoto et al., 2012; Jiang et al., 2012; Nagaya et al., 1987). Numerals indicate positions relative to AB602818. Symbols (β, δ) represent the approximate position of primers used to PCR validate the HBV insertion. The gel image at right shows the 3’ junction PCR results. Ladder band sizes are as indicated. The red filled triangle indicates an on-target product confirmed by capillary sequencing. Repeated attempts to PCR amplify the 5’ junction of the HBV integrant did not return an on-target product, perhaps due to a genomic deletion at the insertion site.

**(C)** Percentages of exonic (black), intronic (grey) and intergenic (blue) genomic alignment breakpoints for ONT reads also aligned to the SARS-CoV-2 genome, and also for the non-reference L1-mediated insertions reported here. Genomic features were annotated according to RefSeq coordinates.

Please see Tables S1 and S2 for further information.

## STAR METHODS

### RESOURCE AVAILABILITY

#### Lead contact

Further information and requests for resources and reagents should be directed to and will be fulfilled by the Lead Contact, Geoffrey J. Faulkner.

#### Materials availability

This study did not generate new unique reagents.

#### Data and code availability

Oxford Nanopore Technologies sequencing data generated by this study were deposited in the European Nucleotide Archive (ENA) under project PRJEB44816. TLDR and instructions for its use and application are available at https://github.com/adamewing/TLDR.

### EXPERIMENTAL MODEL AND SUBJECT DETAILS

Liver tumour and non-tumour tissue were previously obtained from a HBV-positive patient (HCC32, male, 73yrs) who underwent surgical resection at the Centre Hepatobiliaire, Paul-Brousse Hospital, and made available for research purposes with approval from the French Institute of Medical Research and Health (Reference: 11-047). Further ethics approvals were provided by the Mater Health Services Human Research Ethics Committee (Reference: HREC-15-MHS-52) and the University of Queensland Medical Research Review Committee (Reference: 2014000221). HEK293T and Vero E6 cells were obtained from the American Type Culture Collection (ATCC).

## METHOD DETAILS

### SARS-CoV-2 infection of HEK293T cells

HEK293T cells and African green monkey kidney cells (Vero E6) were maintained in standard Dulbecco’s Modified Eagle Medium (DMEM). Culture media were supplemented with sodium pyruvate (11mg/L), penicillin (100U/mL), streptomycin (100μg/mL) (P/S) and 10% foetal calf serum (FCS) (Bovogen, USA). Cells were maintained at 37 °C with 5% CO_2_.

An early Australian SARS-CoV-2 isolate (hCoV-19/Australia/QLD02/2020; GISAID Accession EPI_ISL_407896) was sampled from patient nasopharyngeal aspirates by Queensland Health Forensic and Scientific Services and used to inoculate Vero E6 African green monkey kidney cells (passage 2). A viral stock (passage 3) was then generated on Vero E6 cells and stored at -80°C. Viral titration was determined by immuno-plaque assay (iPA), as previously described (Amarilla et al., 2021). To verify viral replication in HEK293T cells, a growth kinetic was assessed using a multiplicity of infection (MOI) of 0.01, 0.1 or 1.0, and showed efficient SARS-CoV-2 replication (**Figure S1**).

HEK293T viral infection was undertaken as follows: 3×10_6_ HEK293T cells were seeded onto 6-well plates pre-coated with polylysine one day before infection. Cells were infected at MOI of 1 in 200µL of DMEM (2% FCS and P/S) and incubated for 30min at 37°C. Plates were rocked every 5min to ensure the monolayer remained covered with inoculum. The inoculum was then removed, and the monolayer washed five times with 1mL of additive-free DMEM. Finally, cells were maintained with 3mL of DMEM (supplemented with 2% foetal bovine serum and P/S) and incubated at 37°C with 5% CO_2_. Cell supernatant was harvested 0, 1, 2 and 3 days post-infection. The mock infected control differed only in that virus was not added to the inoculum media.

Genomic DNA was extracted from mock and SARS-CoV-2 infected (MOI 1.0) HEK293T cells sampled 2 days post-infection, using a Nanobind CBB Big DNA Kit (Circulomics) following the manufacturer’s instructions for high molecular weight (HMW) DNA extraction. DNA was eluted in elution buffer (10mM Tris-Cl, pH 8.5) and concentration measured by Qubit dsDNA High-Sensitivity Assay Kit on a Qubit Fluorometer (Life Technologies).

### Hepatocellular carcinoma sample processing

DNA was extracted from the HCC32 tissues in our earlier study (Shukla et al., 2013) with a DNeasy Blood and Tissue Kit (QIAGEN, Germany) and stored at -80°C. To enrich for HMW DNA, 4.5μg of DNA from the patient HCC32 tumour and non-tumour liver samples was diluted to 75ng/μL in a 1.5mL Eppendorf DNA LoBind tube and processed with a Short Read Eliminator XS Kit (Circulomics) following the manufacturer’s instructions.

### ONT sequencing

DNA libraries were prepared at the Kinghorn Centre for Clinical Genomics (KCCG) using 3-4μg HMW input DNA, without shearing, and a SQK-LSK110 ligation sequencing kit. 350-500ng of each prepared library was sequenced separately on one PromethION (Oxford Nanopore Technologies) flow cell (FLO-PRO002, R9.4.1 chemistry) (**Table S1**). SARS-CoV-2 infected HEK293T DNA was sequenced on two flow cells. Flow cells were washed (nuclease flush) and reloaded at 24hr and 48hr with 350-500ng of additional library to maximise output. Bases were called with guppy 4.0.11 (Oxford Nanopore Technologies).

### ONT bioinformatic analyses

To call non-reference insertions with TLDR (Ewing et al., 2020), ONT reads generated here, by Zhang *et al*., and by our previous ONT study of human tissues (Ewing et al., 2020) (**Table S1**) were aligned to the human reference genome build hg38 using minimap2 (Li, 2018) version 2.17 (index parameter: -x map-ont; alignment parameters: -ax map-ont -L -t 32) and SAMtools (Li et al., 2009) version 1.12. BAM files were then processed as a group with TLDR (Ewing et al., 2020) version 1.1 (parameters -e virus.fa -p 128 -m 1 --max_te_len 40000 --max_cluster_size 100 --min_te_len 100 --keep_pickles -n nonref.collection.hg38.chr.bed.gz). The file virus.fa was composed of: representative HBV and HCV isolate genomes (**Table S1**), the SARS-CoV-2 isolate used here (GISAID Accession EPI_ISL_407896), the L1HS sequence L1.3 (Dombroski et al., 1993) (Genbank Accession L19088), several *Alu* and SVA subfamily consensus sequences, and a consensus sequence for human endogenous retrovirus K (HERVK), the youngest human long terminal repeat (LTR) retrotransposon family. The file nonref.collection.hg38.chr.bed.gz is a collection of known non-reference retrotransposon insertions available from github.com/adamewing/tldr/. The TLDR output table was further processed to remove calls not passing all TLDR filters, representing homopolymer insertions, where MedianMapQ < 50 or family = “NA” or remappable = “FALSE” or UnmapCover < 0.75 or LengthIns < 100 or EndTE-StartTE < 100 or strand = “None” or SpanReads < 1. As 3’ truncation is rarely encountered for L1-mediated insertions, calls where EndTE was more than 10bp less than the consensus length were filtered, as were *Alu* insertions 5’ truncated by more than 1bp. The filtered TLDR output table is provided as **Table S2**. Insertions detected in only our mock or SARS-CoV-2 infected HEK293T datasets, but not in both experiments, and not matching a known non-reference insertion, were designated as putative cultivar-specific insertions (**Table S2**). Many if not most of these insertions were likely to have occurred in cell culture prior to the cultivars being separated.

To identify L1HS and viral sequences, we directly aligned all reads to the virus.fa file with minimap2 (index parameter: -x map-ont; alignment parameters: -ax map-ont -L -t 32). Reads containing alignments of ≥100bp to a sequence present in virus.fa were counted with SAMtools idxstats. Alignments to HBV, HCV or SARS-CoV-2 were excluded if they overlapped by ≥10bp with a genomic alignment of ≥100bp. Read alignments were visualised with SAMtools view and the Integrative Genomics Viewer (Robinson et al., 2011) version 2.8.6.

### PCR validation

We used Primer3 (Untergasser et al., 2012) to design PCR primers for 6 L1 insertions found by a single spanning ONT read, using the reference genome and L1HS sequences as inputs (**Table S2**). These validation experiments were conducted in three phases. Firstly, we performed an “empty/filled site” PCR using primers positioned on either side of the L1, where the filled site is the L1 allele, and the empty site is the remaining allele(s). Each empty/filled reaction was performed using a DNA Engine Tetrad 2 Thermal Cycler (Bio-Rad) and Expand Long Range Enzyme Mix, with 1X Expand Long Range Buffer with MgCl_2_, 50pmol of each primer, 0.5mM dNTPs, 5% DMSO, 100ng of template DNA and 1.75U of enzyme, in a 25μL final volume. PCR cycling conditions were as follows: (92°C, 3min)×1; (92°C, 30sec; 54-57°C, 30sec; 68°C, 7min)×10; (92°C, 30sec; 52-55°C, 30sec; 68°C, 7min + 20sec/cycle)×30; (68°C, 10min; 4°C, hold)×1. Amplicons were visualised on a 1% agarose gel stained with SYBR SAFE (Invitrogen). GeneRuler_TM_ 1kb plus (Thermo Scientific) was used as the ladder. Secondly, we combined each empty/filled primer with a primer positioned within the L1 sequence, to amplify the 5’ and 3’ L1-genome junctions.

These reactions were undertaken on a DNA Engine Tetrad 2 Thermal Cycler (Bio-Rad), with MyTaq HS DNA polymerase, 1X MyTaq Reaction Buffer, 10pmol of each primer, 10ng of template DNA, and 2.5U of enzyme, in a 25μL final volume. PCR cycling conditions were as follows: (95°C, 1min)×1; (95°C, 15sec; 53-55°C, 15sec; 72°C, 15sec)×35; (72°C, 5min; 4°C, hold)×1. Amplicons were visualised on a 1.5% agarose gel stained with SYBR SAFE (Invitrogen). Thirdly, we repeated the 5’ L1-genome junction-specific PCR using 200ng template DNA. All PCRs were performed with non-template control, as well as DNA extracted from the same HEK293T cells (SARS-CoV-2 and mock) subjected to genomic analysis. Notably, L1 insertions that did not amplify in either cultivar were still likely to be genuine events as they carried all of the relevant sequence hallmarks of L1-mediated retrotransposition.

PCR primers for the HBV insertion 3’ junction (**Figure 3B** and **Table S2**) were designed with Primer3 using the reference genome and closest match HBV sequence (Genbank accession AB602818) as inputs. PCR amplification and capillary sequencing was conducted as per the L1 insertions, except using Expand Long Range polymerase (Roche) with 1X Expand Long Range buffer with MgCl_2_, 10pmol of each primer, 100ng of template DNA, 500µM of PCR Nucleotide Mix, and 3.5U of enzyme, in a 25μL final volume. PCR cycling conditions were as follows: (92°C, 2min)×1; (92°C, 15sec; 65°C, 15sec; 68°C, 7:30min)×10; (92°C, 15sec; 65°C, 15sec; 68°C, 7min+ 20sec per cycle)×35 (68°C, 10min; 4°C, hold)×1. Amplicons were visualized on a 1.2% agarose gel.

Amplicons in each experiment were visualised using a GelDoc (Bio-Rad) and, if of the correct size, gel extracted using a Qiagen MinElute Gel Extraction Kit and capillary sequenced by the Australian Genomics Research Facility (Brisbane).

## QUANTIFICATION AND STATISTICAL ANALYSIS

Error bars and replicate values are defined in figure legends, where appropriate. No statistical tests for significance were conducted.

## KEY RESOURCES TABLE

| REAGENT OR RESOURCE | SOURCE | IDENTIFIER |
| --- | --- | --- |
| **Bacterial and Virus Strains** |  |  |
| SARS-CoV-2 isolate hCoV-19/Australia/QLD02/2020 | Queensland Health Forensic and Scientific Services | EPI_ISL_407896 |
| **Biological Samples** |  |  |
| Snap frozen hepatocellular carcinoma and matched non-tumour liver tissue | Centre Hépatobiliaire, Paul-Brousse Hospital | HCC32 |
| **Chemicals, Peptides, and Recombinant Proteins** |  |  |
| SYBR Safe DNA Gel Stain | Invitrogen | S33102 |
| Agarose | Bioline | BIO-41026 |
| Carboxymethylcellulose | Sigma-Aldrich | C4888 |
| KPL Milk Diluent/Blocking Solution Concentrate | SeraCare | 5140-0011 |
| IRDye 800CW Goat anti-Mouse IgG Antibody | LI-COR | 926-32210 |
| Anti-SARS-CoV-2-spike antibody | Amarilla et al., 2021 | CR3022 |
| **Critical Commercial Assays** |  |  |
| Ligation Sequencing Kit | Oxford Nanopore Technologies | SQK-LSK110 |
| Qubit dsDNA HS Assay Kit | Invitrogen | Q32851 |
| Nanobind CBB Big DNA Kit | Circulomics | NB-900-001-01 |
| Short Read Eliminator XS Kit | Circulomics | SS-100-121-01 |
| Expand Long Template PCR System | Merck | 11681834001 |
| MyTaq DNA Polymerase | Bioline | BIO-21105 |
| **Deposited Data** |  |  |
| Nanopore WGS of mock and SARS-CoV-2 infected HEK293T cells, and HBV infected samples from liver cancer patient HCC32 | This paper | PRJEB44816 |
| Nanopore WGS of SARS-CoV-2 infected HEK293T cells overexpressing L1 | Zhang et al., 2021 | PRJNA721333 |
| Nanopore WGS of normal liver and HCV infected samples from liver cancer patient HCC33 | Ewing et al., 2020 | PRJNA629858 |
| **Experimental Models: Cell Lines** |  |  |
| Human embryonic kidney 293T (HEK293T) cells | ATCC | CRL-1568 |
| *Cercopithecus aethiops* Vero E6 cells | ATCC | CRL-3216 |
| **Oligonucleotides** |  |  |
| Oligo sequences are shown in Table S2. | Integrated DNA Technologies | N/A |
| **Software and Algorithms** |  |  |
| TLDR | https://github.com/adamewing/tldr | Ewing et al., 2020 |
| Minimap2 | https://github.com/lh3/minimap2 | Li et al., 2018 |
| SAMtools | https://github.com/samtools/samtools/ | Li et al., 2009 |
| IGV | https://software.broadinstitute.org/software/igv/ | Robinson et al., 2011 |
| Primer3 | https://bioinfo.ut.ee/primer3-0.4.0/ | Untergasser et al., 2012 |

---

**Figure S1.**
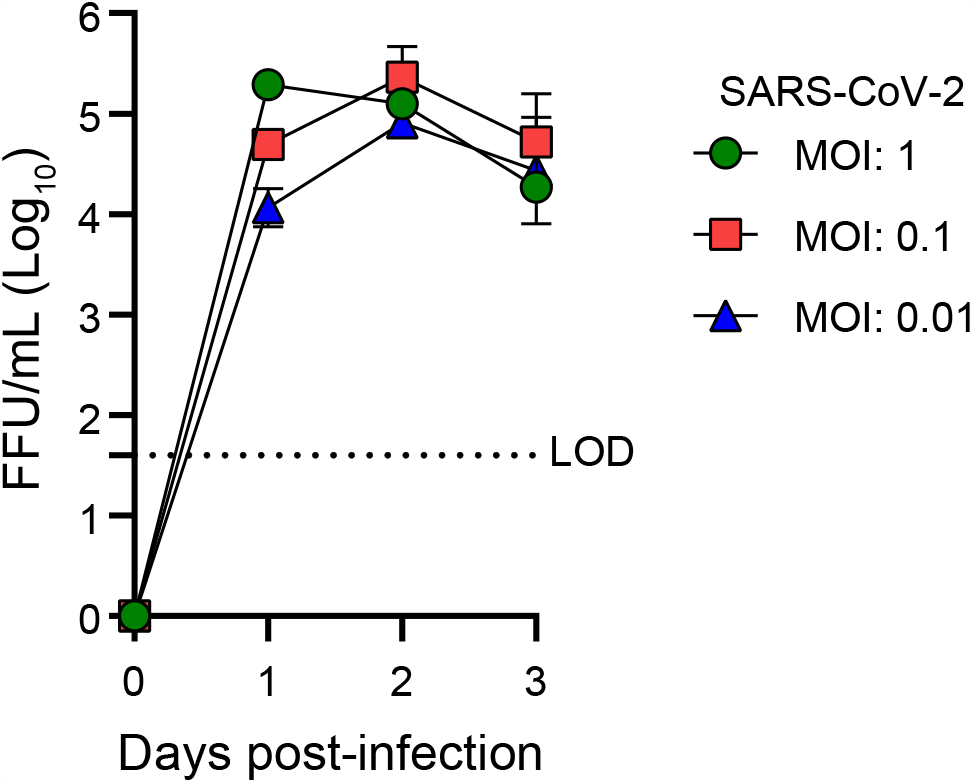
SARS-CoV-2 is replication competent in HEK293T cells, related to Figure 2. HEK293T cells were infected with SARS-CoV-2 isolate QLD02 at an MOI of 0.01, 0.1 and 1.0. Inoculum was removed after infection and cells were washed before the addition of growth media. Supernatant was collected at the indicated time points and viral titres were quantified as focus-forming units (FFU) per mL by immuno-plaque assay (iPA) (Amarilla et al., 2021) with a limit of detection (LOD) as indicated. Data are represented as the mean ± standard deviation of three replicates.

**Figure S2.**
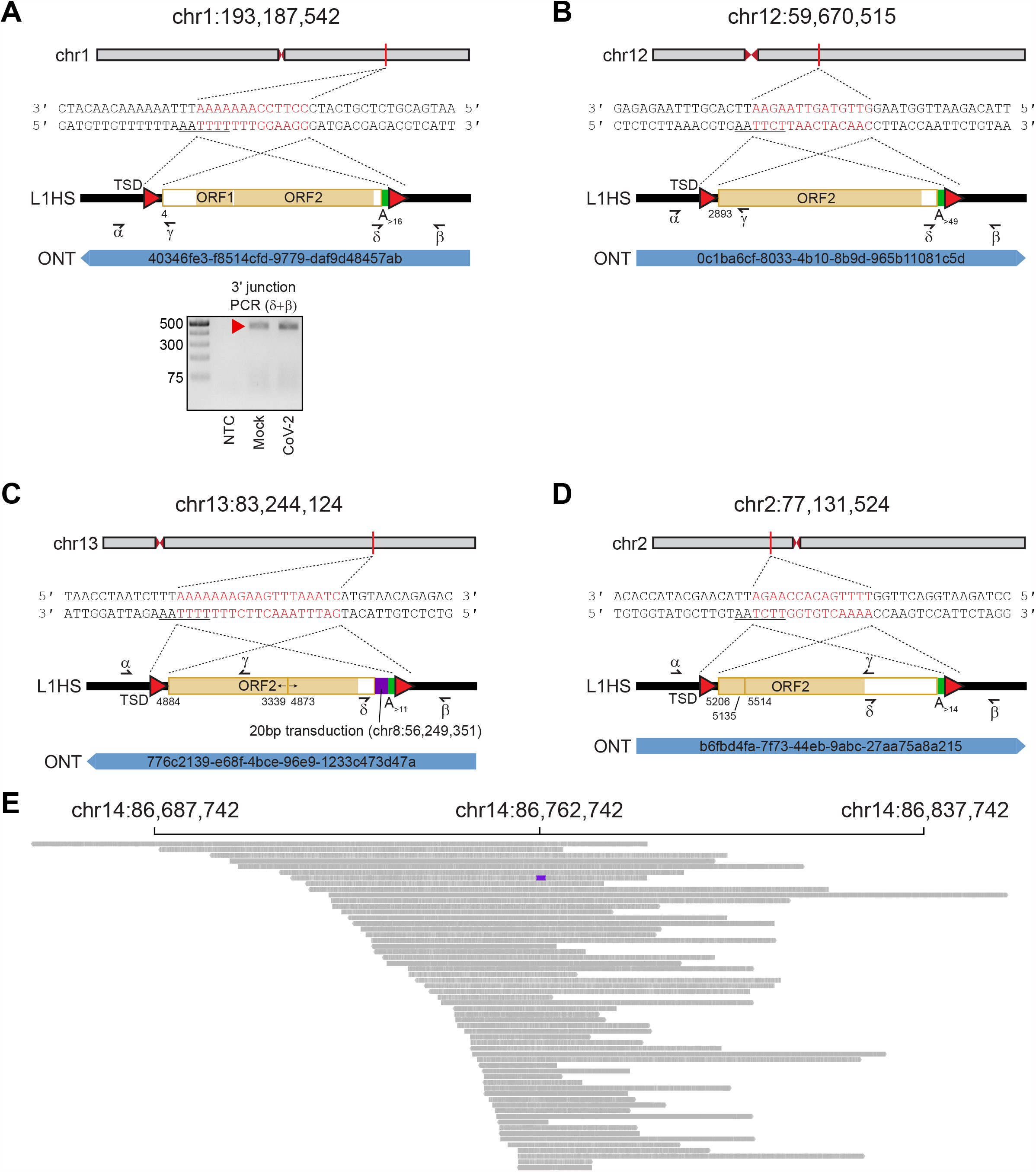
Additional L1HS insertions detected by ONT sequencing in HEK293T cells, related to Figure 2. **(A)** A near full-length L1. **(B)** A 5’ truncated L1. **(C)** A 5’ inverted/deleted L1 carrying a 3’ transduction (purple rectangle) traced to a non-reference source L1. **(D)** A 5’ inverted/deleted L1. **(E)** Integrative Genomics Viewer (Robinson et al., 2011) visualisation of read alignments spanning the L1 integration site displayed in Figure 2D. The L1 is coloured purple. Note: panels (A-D) show the genomic coordinates of an L1 insertion, as well as the sequence at the insertion site. Nucleotides highlighted in red correspond to the integration site TSD. Underlined nucleotides correspond to the L1 EN motif. Cartoons summarise the features of each L1, with the underneath numerals representing the 5’ end position relative to the mobile L1HS sequence L1.3 (Dombroski et al., 1993), TSDs shown as red triangles, and 3’ polyA tracts coloured as green rectangles. One spanning ONT read with its identifier is positioned underneath each cartoon. Symbols (α, β, δ, γ) represent the approximate position of primers used for empty/filled and L1-genome junction PCR validation reactions. The results of the L1-genome 3’ junction PCR are shown for panel (A). Ladder band sizes are as indicated, NTC; non-template control. The red filled triangle indicates an on-target product confirmed by capillary sequencing. No on-target products were observed for the corresponding 5’ junction PCR or the examples shown in panels (B-D).

